# Plasma of COVID-19 patients does not alter electrical resistance of human endothelial blood-brain barrier *in vitro*

**DOI:** 10.1101/2023.09.28.559927

**Authors:** Agnė Pociūtė, Karolina Kriaučiūnaitė, Aida Kaušylė, Birutė Zablockienė, Tadas Alčauskas, Augustė Jelinskaitė, Akvilė Rudėnaitė, Ligita Jančorienė, Saulius Ročka, Alexei Verkhratsky, Augustas Pivoriūnas

**Affiliations:** Department of Stem Cell Biology, State Research Institute Centre for Innovative Medicine, LT-01102, Vilnius, Lithuania; Faculty of Biology, Medicine and Health, The University of Manchester, Manchester, M13 9PT, UK; Achucarro Centre for Neuroscience, IKERBASQUE, Basque Foundation for Science, 48011 Bilbao, Spain; Department of Forensic Analytical Toxicology, School of Forensic Medicine, China Medical University, Shenyang, China; Faculty of Medicine, Vilnius University, LT-03101, Vilnius, Lithuania; Center of Neurosurgery, Vilnius University Hospital Santaros Klinikos, LT-08661, Vilnius, Lithuania; Centre of Infectious Diseases, Vilnius University Hospital Santaros Klinikos, LT-08406, Vilnius, Lithuania

**Keywords:** COVID-19, Blood-brain barrier, TEER, Cytokines, Plasma

## Abstract

The pandemic of Coronavirus Disease 2019 (COVID-19) caused by severe acute respiratory syndrome coronavirus 2 (SARS-CoV-2) instigated the most serious global health crisis. Clinical presentation of COVID-19 frequently includes severe neurological and neuropsychiatric symptoms. However, it is presently unknown whether and to which extent pathological impairment of blood-brain barrier (BBB) contributes to the development of neuropathology during COVID-19 progression.

In the present study we used human induced pluripotent stem cells-derived brain endothelial cells (iBECs) to study the effects of blood plasma derived from COVID-19 patients on the BBB integrity *in vitro*. We also performed a comprehensive analysis of the cytokine and chemokine profiles in the plasma of COVID-19 patients, healthy and recovered individuals.

We found significantly increased levels of interferon γ-induced protein 10 kDa (IP-10), hepatocyte growth factor (HGF), and interleukin-18 (IL-18) in the plasma of COVID-19 patients. However, blood plasma from COVID-19 patients did not affect transendothelial electrical resistance (TEER) in iBEC monolayers.

Our results demonstrate that COVID-19-associated blood plasma inflammatory factors do not impair BBB integrity directly and suggest that pathological remodelling of BBB during COVID-19 may occur through indirect mechanisms.

## Introduction

The COVID-19 pandemic caused by the severe acute respiratory syndrome coronavirus 2 (SARS-CoV-2) has resulted in significant morbidity and mortality worldwide. While primarily targeting the respiratory system, SARS-CoV-2 frequently affects the central nervous system (CNS) by both direct and indirect mechanisms (Carvalho et al., 2021; Steardo et al., 2020a; Steardo et al., 2020b; Tremblay et al., 2020). The COVID-19 infection is associated with a wide range of neurological or neuropsychiatric symptoms, such as anosmia and ageusia, headache, dizziness, confusion, encephalopathy, ischemic stroke, Guillain-Barré syndrome, cognitive impairment, and others (Ellul et al., 2020; Liotta et al., 2020; Mao et al., 2020; Steardo et al., 2020a).

Several *post-mortem* studies have demonstrated the presence of SARS-CoV-2 viral RNA and proteins in various regions of the brain (Matschke et al., 2020; Meinhardt et al., 2021; Thakur et al., 2021) showing direct entry of viral particles into the CNS. Conceptually, there are two possible routes for SARS-CoV-2 virus invasion into the brain. First, SARS-CoV-2 can infect the sustentacular glial cells of the olfactory epithelium in the nasal cavity, subsequently gaining access to the brain through the olfactory nerve (Butowt and von Bartheld, 2021; Coolen et al., 2020; Verma et al., 2022). Second, SARS-CoV-2 virus can gain an access to the brain parenchyma through disrupted blood-brain barrier (BBB). The BBB is a highly selective and dynamic interface that separates the blood from the CNS (Pivoriunas and Verkhratsky, 2021; Verkhratsky and Pivoriunas, 2023). It is composed of specialised brain endothelial cells (BECs) that line the brain vessels, along with astrocytes, pericytes, parenchymal and endothelial basement membranes and perivascular space that mount the barrier protecting the CNS from pathogens and toxins (Sweeney et al., 2019; Verkhratsky and Butt, 2023). Accumulating evidence indicate that cerebral vascular dysfunction is a common feature of COVID-19: *post-mortem* studies on COVID-19 patients have shown haemorrhages and ischaemia in the CNS (Jensen et al., 2021; Matschke et al., 2020; Thakur et al., 2021), whereas imaging studies on recovering patients similarly revealed abnormalities in cerebral blood flow and hypometabolism, indicating disruption of the BBB (Vanderheiden and Klein, 2022).

The SARS-CoV-2 virus enters host cells through the angiotensin-converting enzyme 2 (ACE2) receptor, which is expressed in various tissues, including BECs (Buzhdygan et al., 2020) and therefore can target BBB directly. It was shown that SARS-CoV-2 virus infects and replicates in cultured human inducible pluripotent stem cell (iPSC)-derived BECs (iBECs), which however does not affect barrier function (Krasemann et al., 2022). At the same time, the SARS-CoV-2 spike proteins directly interact with BECs *in vitro* and may alter BBB function (Motta et al., 2023; Vanderheiden and Klein, 2022). However, infection of BECs with SARS-CoV-2 *in vivo* has not yet been demonstrated.

COVID-19 is accompanied by an excessive systemic release of pro-inflammatory cytokines into the blood, a phenomenon known as cytokine storm (Montazersaheb et al., 2022). Several specific cytokine and chemokine profiles have been linked to COVID-19 severity and neurological complications (Ye et al., 2020; Mehta et al., 2020) (Mehta et al., 2020; Ye et al., 2020). However, the effects of cytokines and chemokines secreted during SARS-CoV-2 infection on the BBB function were not systematically investigated. Pro-inflammatory cytokines and chemokines can affect BBB by following mechanisms: (i) directly affecting BECs thus causing disruption of the BBB; (ii) stimulating release of inflammatory mediators by BECs into the brain parenchyma to induce reactive astrogliosis and microgliosis leading to the secondary BBB damage; (iii) pro-inflammatory cytokines and chemokines can penetrate BBB causing neuroinflammation leading to the secondary BBB injuries; (iv) combinations of above.

Therefore, it is important to understand the intricate connections between blood cytokine/chemokine profiles, BBB integrity and the severity of neurological manifestations of COVID-19. This may facilitate the development of potent diagnostic tools enabling early prediction and prevention of neurologic complications in COVID-19 patients.

Current technologies enable the generation of human iBECs monocultures with transendothelial electrical resistance (TEER) in the range of 4000-5000 Ω.cm^2^ that is close to the readings obtained *in vivo* (Kriauciunaite et al., 2021; Neal et al., 2019). In the present study, we directly monitored BBB function in response to the blood plasma from COVID-19 patients. We also characterised blood plasma cytokine/chemokine profiles and found significantly increased levels of interferon γ-induced protein 10 kDa (IP-10), hepatocyte growth factor (HGF), and interleukin-18 (IL-18) in the plasma of COVID-19 patients when compared to healthy and/or recovered individuals. However, blood plasma from COVID-19 patients did not affect BBB electrical resistance. Our results show that COVID-19-associated blood plasma inflammatory factors do not damage BBB directly and suggest that pathological remodelling (if any) of BBB during COVID-19 may occur through indirect mechanisms.

## Methods

### Patient information and data collection

This study examined 33 patients who were diagnosed with COVID-19 and were admitted to the Centre of Infectious Diseases of Vilnius University Hospital Santaros Klinikos between 2022 and 2023. We also assessed 30 healthy and 18 recovered subjects. Clinical information and laboratory samples were collected immediately after hospitalisation. The permission to conduct this biomedical research was issued by the Vilnius Regional Biomedical Research Ethics Committee (2022/2-1407-879). The median age of the subjects was 46 years (IQR 38 – 58). The youngest subject was 20 and the oldest was 65 years old, 37 (46.27%) of the subjects were female and 43 (53.75%) were male.

All study participants voluntarily gave informed consent to participate in the study.

### Collection of plasma samples

Whole blood samples were collected in ethylenediaminetetraacetic acid (EDTA) and sodium heparin tubes. Within 30 min. samples were centrifuged at 1000 × g for 10 min at 4 °C to obtain plasma. Plasma was transferred in conical centrifuge tubes and subjected to inactivation at 56 °C for 30 min. Afterwards, plasma was centrifuged at 1000 ×g for 10 min at 4 °C, aliquoted in cryotubes, and stored at -80 °C until further use.

### Maintenance of iPSCs and differentiation to the brain capillary endothelial cells (iBECs)

Human exfoliated deciduous teeth stem cell (SHED)-derived iPSCs (female, 7 years old) were cultured on matrigel-coated (Corning) plates with Essential 8 Flex medium (E8) in the incubator (37°C and 5% CO^2^). iPSCs were differentiated to iBECs according to slightly modified previously published protocol (Neal et al., 2019). Briefly, 24 hours after splitting, the differentiation was initiated by changing the E8 to the Essential 6 (E6) medium (Thermo Fisher Scientific). E6 was fully refreshed every 24 hours for four days. On the fifth day, the E6 was changed to the human Endothelial Serum-Free Medium (hESFM, Thermo Fisher Scientific) supplemented with 20 ng/ml bFGF (Thermo Fisher Scientific), 10 μM retinoic acid (Merck Darmstadt, Germany), and 0.25× B-27 (Thermo Fisher Scientific). After 48 hours the same medium was fully refreshed. The next day cells were split for selection on 400 μg/ml collagen-IV and 100 μg/ml fibronectin-coated (both from Merck) Transwell inserts (Corning) in hESFM medium supplemented with 0.25× B-27, 50 U/ml and 50 µg/ml penicillin-streptomycin (Thermo Fisher Scientific, hESFM+B-27+P/S).

### Measurement of transendothelial electrical resistance

TEER monitoring was used as a readout of BBB integrity. TEER of iBECs monolayers was monitored every hour using the CellZscope system (NanoAnalytics, Münster, Germany), beginning immediately after seeding cells on the insert (at -24 h) and were carried out for 24-48 hours in the presence of plasma after its addition at 0-hour time point. The inserts plated with iBECs were immediately loaded into a 24-well cell module of a CellZscope system prefilled with hESFM+B-27+P/S and grown in the incubator (37°C and 5% CO^2^). 24 hours after plating, the hESFM+B-27+P/S was refreshed, and the cells were treated with 50 % plasma (v/v in the same medium) collected from three different groups: healthy volunteers, COVID-19 infected patients, and recovered COVID-19 patients.

### Immunofluorescence

After TEER measurements, the culture medium was aspirated and the cells were washed with PBS 3 times. Cells were fixed and permeabilized using ice-cold (-20 °C) methanol-acetone solution (1:1), for 10 min at -20 °C. Then the membrane of the Transwell insert was cut out and placed on the parafilm. Cells were washed 3 times with PBS, blocked using 1% bovine serum albumin-PBS solution for 30 min at room temperature, then incubated with primary antibodies (against ZO-1 (1:33), occludin (1:50), and claudin-5 (1:100), in 1% bovine serum albumin-PBS) overnight at 4 °C. After incubation, samples were washed 3 times with PBS and incubated with anti-rabbit IgG secondary antibodies conjugated with an Alexa Fluor-594, diluted in PBS (1:1000) for 1 hour at room temperature in the dark. After incubation, the cells were washed again 3 times with PBS. Membrane with cells were placed on the objective slide in mounting medium with DAPI and covered with a coverslip. Samples were visualized with a Leica TCS SP8 confocal microscope using Diode 405 nm, DPSS 561 nm, lasers. Photos were taken with a 63x immersion lens.

### Cytokine measurements

For multiplex quantitative cytokine analysis, ProcartaPlex™ immunoassays were applied, using the Luminex 200™ detection system (Invitrogen). Plasma cytokine levels were determined by commercial multiplex immunoassays (Invitrogen, Thermo Fischer Scientific) according to the manufacturer’s instructions. Cytokine concentrations were measured in duplicates from previously unthawed plasma samples using the ProcartaPlex™ Human Cytokine/Chemokine/Growth Factor Convenience Panel 1 45-Plex (EPXR450-12171-901), the ProcartaPlex™ Human MixMatch 17-plex (PPX-17-MXYMK7R), and the ProcartaPlex™ Human RANTES (CCL5) Simplex/Basic Kit (EPX010-10420-901, EPX01A-10287-901).Frozen plasma samples were thawed at room temperature, spun at 1000 g for 10 min, and transferred to a new Eppendorf tube before immunoassay.

The multiplex immunoassays applied in this study consist of most relevant cytokines, chemokines, and growth factors typically found in inflammation. The following cytokines were analysed: brain-derived neurotrophic factor (BDNF), epidermal growth factor (EGF), eotaxin (CCL11), fibroblast growth factor (FGF-2), granulocyte–macrophage colony stimulating factor (GM-CSF), growth-regulated protein-α (GRO-α /CXCL1), hepatocyte growth factor (HGF), interferon-α (IFN-α), IFN-γ, interleukin 1 receptor antagonist (IL-1RA), IL-1α, IL-1β, IL-2, IL-4, IL-5, IL-6, IL-7, IL-8, IL-9, IL-10, IL-12p70, IL-13, IL-15, IL-17A (CTLA-18), IL-18, IL-21, IL-22, IL-23, IL-27, IL-31, IFN-γ-inducible protein 10 (IP-10/CXCL10), leukaemia inhibitory factor (LIF), monocyte chemotactic protein 1 (MCP-1/CCL2), macrophage inflammatory protein type 1 α (MIP-1α /CCL3), MIP-1β (CCL4), β-nerve growth factor (NGF-β), regulated on activation, normal T-cell expressed and secreted (RANTES/CCL5), platelet-derived growth factor-BB (PDGF-BB), placenta growth factor 1 (PIGF-1), stem cell factor (SCF), stromal cell-derived factor 1 (SDF-1α), tumour necrosis factor-α (TNF-α), TNF-β, vascular endothelial growth factor-A (VEGF-A), and VEGF-D.

Cytokine concentrations were calculated using the standard curve generated by five-parameter logistic regression method.

### Statistical analysis

Statistical analysis was performed using Graph Pad Prism® software 8.0.2 (Graph Pad Software, Inc., City, State, USA). Differences between the groups were compared by 2way ANOVA (following Tukey’s multiple comparisons test) or non-parametric Kruskal-Wallis test (following Dunn’s multiple comparisons test). Results were considered as significant when *p* < 0.05.

## Results

### Plasma of COVID-19 patients does not affect TEER

To investigate the potential effect of patient-derived blood plasma on the integrity of the BBB, iBECs were seeded on Transwell inserts and treated with patient-derived blood plasma, obtained from three different groups: healthy volunteers, COVID-19 patients, and recovered COVID-19 patients. The treatment involved applying 50% (v/v) plasma to the cells.

We first tested the effects of exposure of endothelial monolayer to pooled plasma administered at the apical, basolateral or both sides of the Transwell inserts. Our results show that plasma contacting only basolateral or apical and basolateral sides completely inhibited TEER of iBEC monolayers, whereas iBECs exposed to the plasma only from the apical side demonstrated normal TEER readings (Fig. 1A).

**Figure 1.**
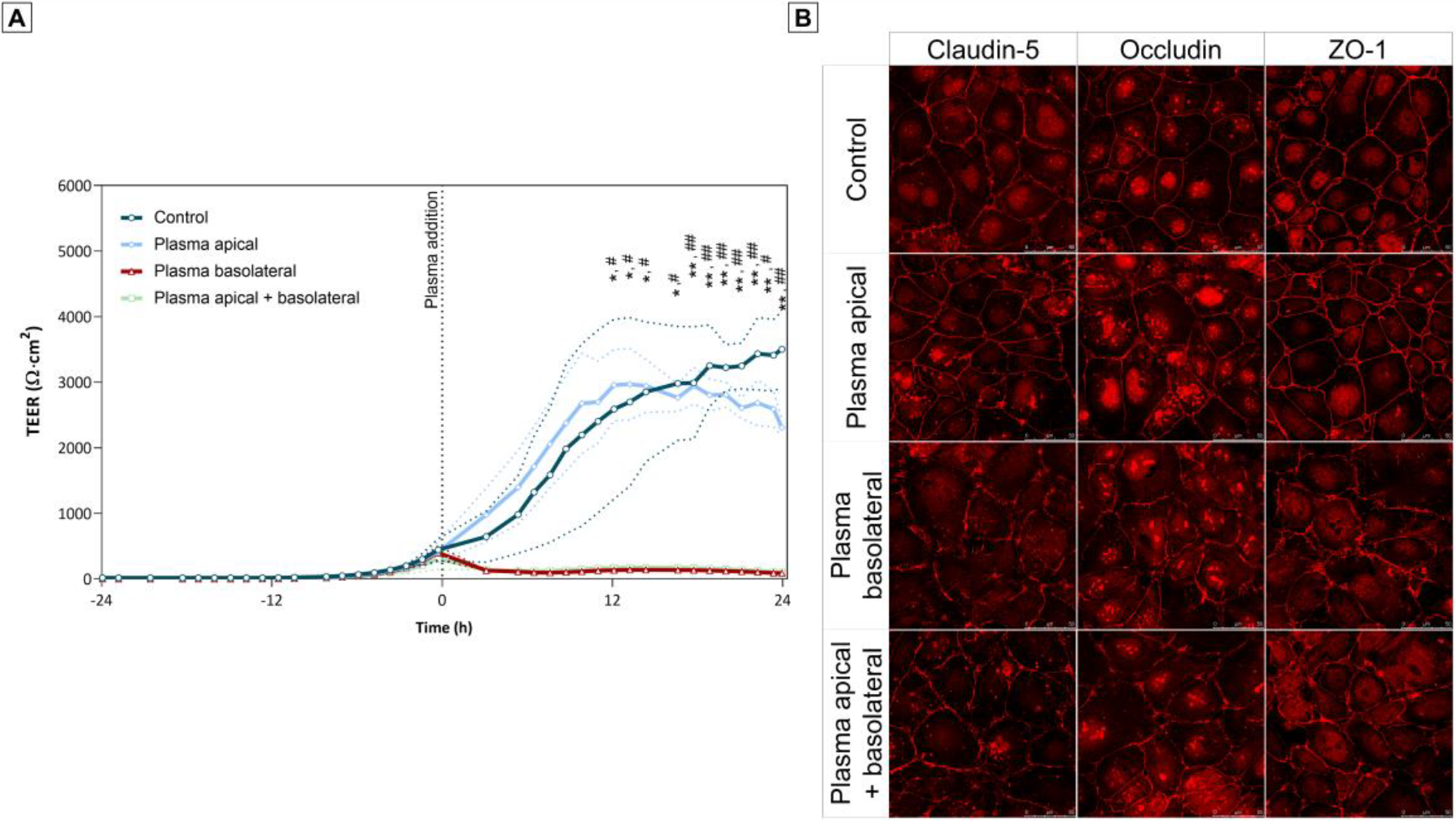
The effects of plasma on human induced brain cerebrovascular endothelial cells (iBECs) exposed from the apical, basolateral or both sides. Cells were treated with 50 % plasma (v/v) from apical, basolateral or both sides of the insert, control – plasma non-treated cells. **(A)** Dynamic changes in transendothelial electrical resistance (TEER). Data are presented as a mean ± S.D. * – Statistically significant difference between apical and basolateral treatment, # – statistically significant difference between apical and apical + basolateral treatment. *, #p < 0.05, **, ##p < 0.01, N = 3. **(B)** Immunofluorescence analysis of iBEC cultures, stained with antibodies against tight junction (TJ) proteins claudin-5, occludin, zonula occludens-1 (ZO-1).

Immediately after TEER measurements, cells were fixed for immunostaining with antibodies against TJ proteins claudin-5, occludin and zonula occludens-1 (ZO-1). Confocal microscopy images showed that exposure to plasma from basolateral and both sides disrupted TJs in iBEC monolayers as evidenced by more irregular and fragmented staining patterns of TJ proteins when compared to controls or cells treated from the apical side (Fig. 1B).

Next, we investigated the effects of plasma from healthy, recovered subjects and COVID-19 patients administered to the apical side of the monolayer on its electrical resistance. Dynamic changes in TEER values show a similar trend for all groups (Fig. 2B), starting to rise around 12 hours after seeding the cells on Transwell inserts and reaching the values of ∼2500 Ω.cm^2^ after 24 h after seeding when the plasma was added (zero time point). Immediately after the addition of plasma, TEER dropped slightly probably reflecting injection artefact. Following this drop, TEER continued to rise steadily and reached its peak exceeding 4000 Ω.cm^2^ around 12 hours after the addition of plasma. After reaching the peak, the TEER slowly started to decline but remained above 2000 Ω.cm^2^ at the end of the 48-hour measurement period. Analysis of TEER data did not reveal any significant differences between experimental groups at any time point during the 48-hour observation period after plasma addition.

**Figure 2.**
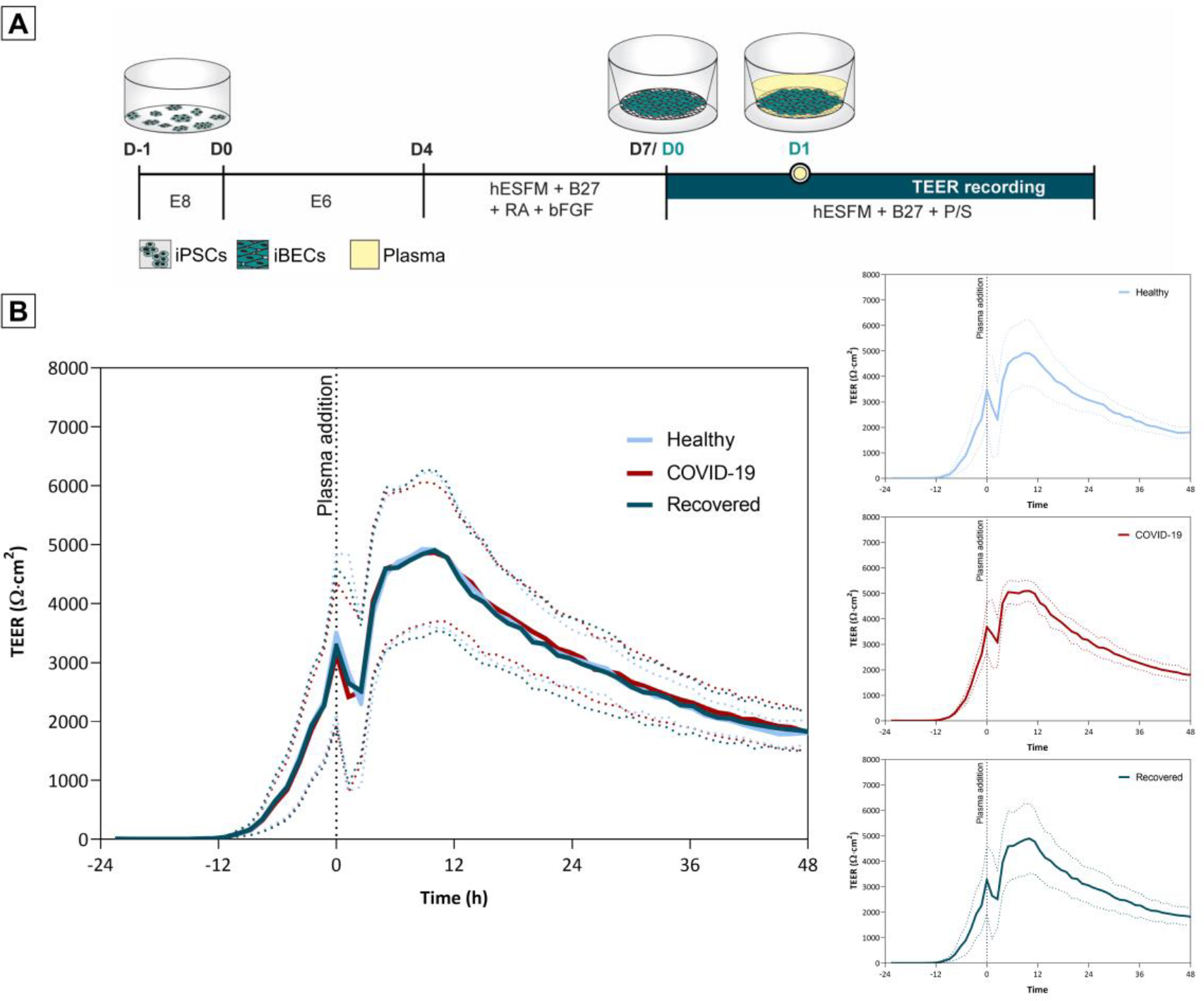
The effect of plasma from healthy volunteers, COVID-19 patients and recovered subjects on transendothelial electrical resistance (TEER) of human induced brain cerebrovascular endothelial cells (iBECs) monolayers. **(A)** Experimental protocol. **(B)** Time course of TEER (Ω·cm^2^) changes of iBECs grown on transwell inserts and treated with plasma from apical side; data are presented as a mean ± S.D. Healthy volunteers (N = 20), COVID-19 patients (N = 24), recovered subjects (N = 17).

### Cytokine profiles of plasma of COVID-19 patients, recovered patients and healthy controls

We compared concentrations of cytokines, chemokines, and growth factors in the plasma of hospitalised COVID-19 patients with healthy and recovered individuals. We first used a 45-analyte multiplex immunoassay panel to measure plasma levels of 45 cytokines in COVID-19 patients (N=18), recovered (N=13), and healthy (N=9) subjects (Fig. 2). Analysis of cytokine profiles showed increased plasma levels of IP-10, HGF, VEGF-A, IL-7, IL-18, and MCP-1/CCL2 in COVID-19 patients. However, statistically significant differences between tested groups were found only for IP-10 and HGF. Concentrations of IP-10 were statistically higher in blood plasma of COVID-19 patients compared to recovered subjects (p<0.05). Statistically significantly higher levels of HGF were detected in the plasma of COVID-19 patients compared to healthy (p<0.05) and recovered (p<0.001) subjects. We also observed that SCF levels in COVID-19 patients were statistically significantly lower than in healthy individuals (p<0.05). eotaxin/CCL11, BDNF, LIF, PIGF-1, PDGF-BB, MIP-1α/CCL3, MIP-1 β/CCL4, RANTES/CCL5, SDF-1α, and VEGF-D were detected in the plasma of all three groups but showed no significant differences between the tested groups. 15 of the 45 analytes assessed were below the lower limit of quantification in 90% of all samples (irrespective of group) and were therefore excluded from further analysis. 12 analytes were observed at detectable levels in some samples of all three groups but showed no significant differences between the tested groups and were also excluded from further analysis.

Therefore, we further determined the levels of 18 cytokines (IP-10, HGF, IL-7, IL-18, IL-31, MCP-1/CCL2, eotaxin/CCL11, BDNF, LIF, PIGF-1, PDGF-BB, MIP-1α/CCL3, MIP-1 β/CCL4, RANTES/CCL5, SDF-1α, SCF, VEGF-A, and VEGF-D) in the plasma of COVID-19 patients (N=13), healthy (N=20), and recovered (N=5) subjects.

The final analysis of the cytokine assays showed that the levels of IP-10, HGF, IL-18, eotaxin, RANTES, and MIP-1β were statistically significantly different between the tested groups (Fig. 3). Concentrations of IP-10 and HGF were statistically higher in blood plasma of COVID-19 patients compared to recovered (p<0.0001) and healthy (p<0.05 and p<0.001) subjects. Statistically significantly higher levels of IL-18 were detected in the plasma of COVID-19 patients compared to recovered subjects (p<0.05). Meanwhile, the levels of eotaxin in healthy subjects were statistically significantly higher than in COVID-19 patients and recovered individuals (p<0.05). Plasma MIP-1β levels were significantly higher in healthy subjects than in recovered individuals (p<0.05), while RANTES levels were significantly higher in recovered than in healthy subjects (p<0.05).

**Figure 3.**
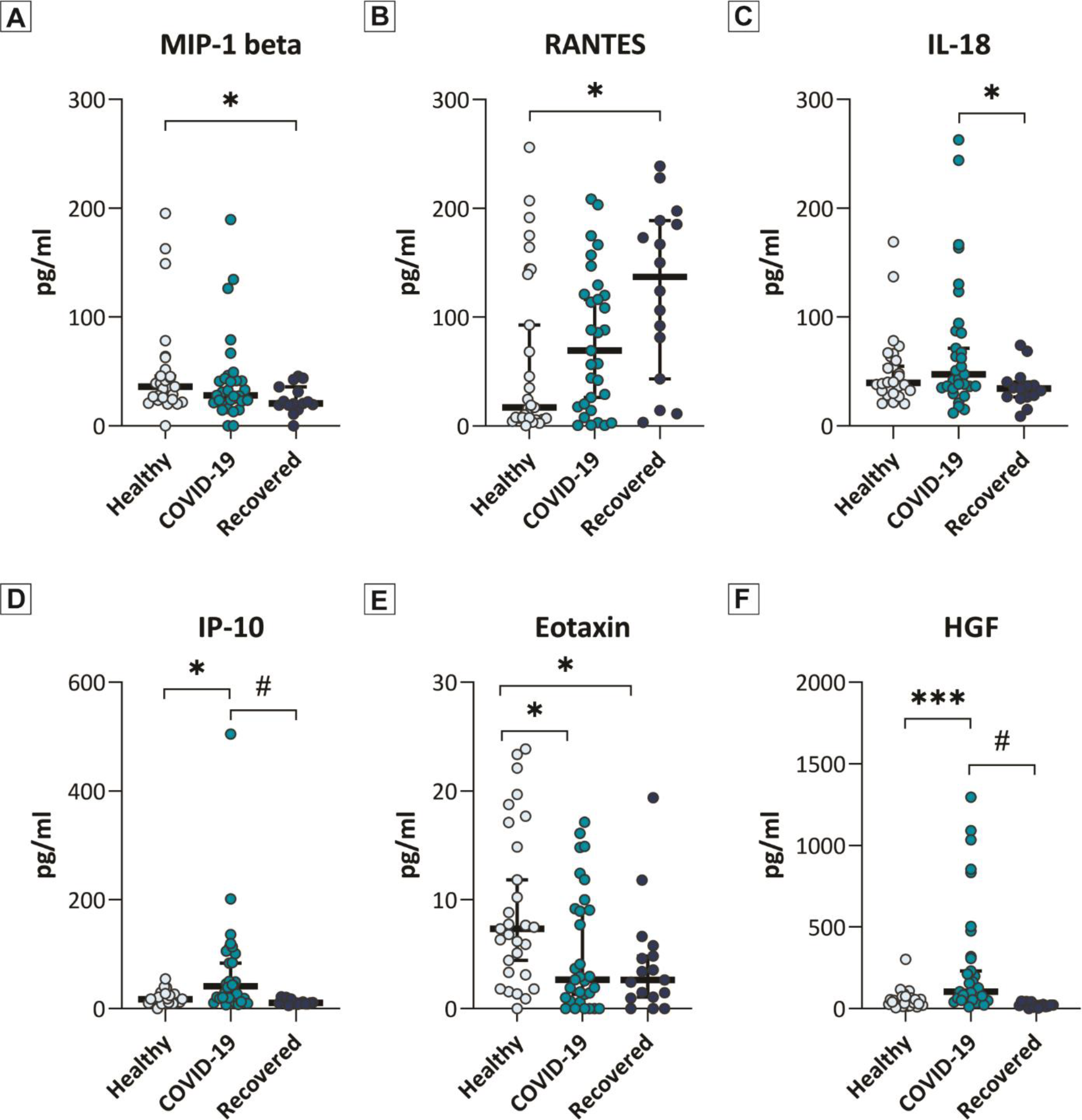
Plasma cytokine profiles of COVID-19 patients, recovered subjects and healthy controls. The plasma cytokine levels of **(A)** MIP-1 beta, **(B)** RANTES, **(C)** IL-18, **(D)** IP-10, **(E)** Eotaxin, and **(F)** HGF from healthy volunteers (N = 29), COVID-19 patients (N = 31) and recovered (N = 18) subjects. Data are presented as a median with 95 % confidence interval, *p < 0.05, ***p < 0.001, #p <0.0001.

Thus, our results showed that COVID-19 significantly increases the levels of IP-10, HGF, and IL-18 in the plasma of COVID-19 patients compared to healthy and/or recovered individuals. As all COVID-19 patients in our study were hospitalised, it is reasonable to consider IP-10, HGF and IL-18 as biomarkers of disease severity in SARS-CoV-2 infected patients.

## Discussion

In the present study we investigated the effects of COVID-19 patient-derived blood plasma on the *in vitro* model of human BBB. The apical side of iBECs monolayers was directly exposed to the heat-inactivated patient or healthy subjects-derived plasma and TEER was continuously monitored during a 48-hour period. Using this experimental model, we were able to achieve peak TEER values exceeding 4000 - 5000 Ω.cm^2^ thus showing compatibility of BECs to the exposure with human plasma. We also found that exposure of iBECs to the plasma either from basolateral or basolateral and apical sides completely disrupted barrier function; at the same time plasma administration at the apical side only did not affect TEER (Fig. 1). Apicobasal polarisation is crucial for the proper barrier function of the BECs (Dreifuss, 1989). Extravasated blood plasma components are directly toxic to the brain parenchyma and also trigger massive secondary disruption of the BBB leading to the brain oedema and tissue necrosis (Aronowski and Zhao, 2011). Our *in vitro* model demonstrates the importance of apicobasal polarity for BEC barrier function and could be used for modelling of different pathological conditions related to the secondary BBB injury.

We found no changes in the TEER values between the experimental groups consisting of SARS-CoV-2-infected patients and healthy or recovered individuals. We suggest that our results could be explained by two groups of reasons: (i) experimental design; (ii) heat-insensitive COVID-19-associated blood plasma inflammatory factors do not affect BEC barrier functions directly.

In this study, we used heat-inactivated blood plasma (56 °C for 30 min). Heat inactivation was necessary to prevent complement-mediated lysis in cell cultures and as a safety precaution for the effective inactivation of SARS-CoV-2 viral particles. However, heat inactivation can significantly affect quantities and biological activity of soluble factors present in plasma. Comparison of proteomic profiles revealed downregulation of many cytokines, chemokines and growth factors in heat-inactivated plasma (Ayache et al., 2006). It was also demonstrated that plasma from patients hospitalised with acute SARS-CoV-2 infection decreased transendothelial resistance of human lung microvascular endothelial cells, but these destructive effects were susceptible to heat inactivation (Kovacs-Kasa et al., 2022). Finally, complement components can by themselves damage BBB. Various injuries may initiate the inflammatory response of BECs by promoting activation and binding of circulating complement components to the abluminal membranes, stimulating the secretion of pro-inflammatory factors and instigating BBB leakage (Orsini et al., 2014; Propson et al., 2021). We, therefore, cannot exclude the possibility that heat-sensitive components present in the plasma of SARS-CoV-2-infected patients may affect the electrical resistance of iBEC monolayers. Further studies are needed to address this question.

Our data demonstrate that heat-insensitive COVID-19-associated blood plasma inflammatory factors do not affect BEC barrier functions directly. However, we do not exclude the possibility that COVID-19-associated plasma factors can trigger BECs to release pro-inflammatory factors into the brain parenchyma leading to secondary BBB damage. For instance, infection of human iPSCs-derived BECs with SARS-CoV-2 did not affect barrier properties but upregulated IFN-γ signalling, and these results were consistent with histopathological studies showing upregulated IFN-γ pathway in COVID-19 human neurovascular unit (Krasemann et al., 2022). Finally, BBB can selectively transport several pro-inflammatory cytokines from the peripheral circulation into the brain parenchyma and promote secondary BBB injury (Pan et al., 2011). In the future, a systematic comparison of molecular responses between BECs exposed to COVID-19 and control plasma may resolve this issue.

After analysing 45 cytokines in the plasma of COVID-19 patients, we found that only the levels of IP-10, HGF, and IL-18 were significantly higher in COVID-19 patients compared to healthy and/or recovered subjects. IL-18 is known as a pro-inflammatory cytokine involved in host defence against infections and regulating innate and acquired immune responses (Ihim et al., 2022). Our results are in agreement with other studies showing correlations of IL-18 serum levels with other markers of inflammation and disease severity (Rosen et al., 2022; Satis et al., 2021; Trifonova et al., 2023).

Interferon-γ-induced protein (IP-10), also known as small inducible cytokine B10, belongs to the CXC chemokine family (also known as CXCL10). This 10-kDa cytokine recruits immune cells, including T cells, natural killer cells, and macrophages, to the inflamed tissue in inflammatory diseases. IP-10 induces T cells and is therefore important for antiviral defence. It is upregulated in the blood of hospitalised COVID-19 patients (Huang et al., 2020). These data corroborate our study, where IP-10 was significantly higher in the plasma of COVID-19 patients than in healthy (p<0.05) and recovered (p<0.0001) subjects. Comparison of cytokine expression profiles between critically ill, severe, and moderate COVID-19 cases revealed a significant association of IP-10 with disease severity (Yang et al., 2020). Previous studies demonstrated that IP-10 is one of the most abundant and the earliest chemokines associated with BBB damage in various viral infections (Wang et al., 2018). Besides COVID-19, IP-10 was also associated with the severity of diseases caused by other viruses such as MERS-CoV and influenza (Faure et al., 2014; Wang et al., 2010). IP-10 is considered as a biomarker of multiple CNS diseases and closely correlates with BBB pathological changes (Amin et al., 2009; Chai et al., 2015; Li et al., 2015). In Japanese encephalitis, for example, IP-10 promoted BBB damage by inducing TNF-α production through the JNK-c-Jun signalling pathway; in turn, TNF-α affected the expression and distribution of tight junctions in brain microvascular endothelial cells resulting in BBB damage (Wang et al., 2018). However, in our study, TNF-α levels were below the lower limit of quantification in 95% of all tested samples. This possibly reflects an inappropriate inflammatory response that may occur during COVID-19 (Gomez Marti et al., 2021).

We also found that plasma HGF levels were significantly higher in COVID-19 patients than in recovered (p<0.001) and healthy (p<0.05) subjects. HGF produced by stromal and mesenchymal cells regulates epithelial cell proliferation, motility, morphogenesis, and angiogenesis (Gupta et al., 2022). HGF is a pleiotropic cytokine with anti-inflammatory properties that plays a key role in lung tissue repair and can modulate the adaptive immune response (Perreau et al., 2021; Sanchez-de Prada et al., 2022). Increased HGF production induced by lung injury promotes tissue repair (Gupta et al., 2022). In general, increased circulation of growth factors such as HGF is associated with repair mechanisms following acute lung injury during SARS-CoV-2 infection (Young et al., 2021). However, a marked increase in HGF was also significantly correlated with disease severity (Gupta et al., 2022; Young et al., 2021). HGF is also a marker of neutrophil activation, it was considered as one of the strongest indicators of critical illness in COVID-19 (Meizlish et al., 2021). Moreover, upregulated HGF in intensive care unit patients, is elevated in chronic inflammatory diseases (Haljasmagi et al., 2020). In our study, HGF levels in the blood of COVID-19 patients were 11 times higher than in recovered subjects (p<0.001) and 5 times higher than in healthy subjects (p<0.05). Our findings are in agreement with studies showing an association of HGF with the severity of COVID-19 (Yang et al., 2020; Perreau et al., 2021).

Throughout the course of the COVID-19 pandemic, various strains of the SARS-CoV-2 virus have emerged, exhibiting different characteristics and potential pathological effects (Markov et al., 2023). It is worth noting that most of the published research investigating cytokine profiles and viral impact on the BBB was conducted during the early and middle stages of the pandemic. However, for our study blood plasma samples were collected during the period from February 2022 to April 2023, which represents a later stage of the pandemic. It is plausible to speculate that the novel circulating viral strains, which emerged during this period, may have undergone genetic variations and evolutionary changes, potentially leading to alterations in cytokine profiles and their influence on BBB.

In conclusion, our study demonstrates that blood plasma from COVID-19 patients, although enriched with pathologically relevant cytokines and chemokines, does not affect BBB electrical resistance. Our findings warrant further research to explore possible indirect mechanisms of pathological BBB remodelling during COVID-19.

## Acknowledgements

This study was supported by funding from European Regional Development Fund (project No 13.1.1-LMT-K-718-05-0005) under grant agreement with the Research Council of Lithuania (LMTLT). Funded as European Union’s measure in response to Cov-19 pandemic.

